# Reconstitution of a minimal ESX-5 type VII secretion system suggests a role for PPE proteins in outer membrane transport of proteins

**DOI:** 10.1101/2022.09.05.506643

**Authors:** C. M. Bunduc, Y. Ding, C. Kuijl, T. C. Marlovits, W. Bitter, E.N.G. Houben

**Author notes:** Corresponding author. Vrije Universiteit Amsterdam, Section Molecular Microbiology, de Boelelaan 1108, 1081 HZ Amsterdam, the Netherlands.

## Abstract

Mycobacteria utilize type VII secretion systems (T7SSs) to secrete proteins across their highly hydrophobic and diderm cell envelope. Pathogenic mycobacteria have up to five different T7SSs, called ESX-1 to ESX-5, which are crucial for growth and virulence. Here, we use a functionally reconstituted ESX-5 system in the avirulent species *Mycobacterium smegmatis* that lacks ESX-5, to define the role of each *esx-5* gene in system functionality. By creating an array of gene deletions and assessing protein levels of components and membrane complex assembly, we observed that only the five components of the inner membrane complex are required for its assembly. However, in addition to these five core components, active secretion also depends on both the Esx and PE/PPE substrates. Tagging the PPE substrates followed by subcellular fractionation, surface labeling and membrane extraction showed that these proteins localize to the mycobacterial outer membrane. This indicates that they could play a role in secretion across this enigmatic outer barrier. These results provide a first full overview of the role of each *esx-5* gene in T7SS functionality.

## Introduction

Gram-negative bacteria have evolved an exceptional variety of specialized secretion systems to mediate protein transport across their diderm cell envelope (Costa *et al*, 2015). Bacterial pathogens usually strictly depend on these systems to complete their infection cycle. Mycobacteria have an unusual diderm cell envelope with a distinctive outer membrane, also known as the mycomembrane, which is mainly composed of mycolic acids (Dulberger *et al*, 2020). These bacteria use a specific group of homologous specialized secretion systems, called type VII secretion systems, for a diverse array of functions, such as intracellular survival, immunomodulation, as well as uptake of nutrients and metabolites (Ates *et al*, 2015; Bunduc *et al*, 2020a; Pym *et al*, 2002; Siegrist *et al*, 2009; Simeone *et al*, 2012). Pathogenic mycobacteria, such as *Mycobacterium tuberculosis*, encode up to five different T7SSs, named ESX-1 to ESX-5 (Bunduc *et al*., 2020a). The importance of T7SS for pathogenicity is further evidenced by the lack of ESX-1 causing the attenuation of the currently used live vaccine strain *Mycobacterium bovis* BCG (Houben *et al*, 2012a; Pym *et al*., 2002; Simeone *et al*., 2012; van der Wel *et al*, 2007).

The mycobacterial *esx* loci contain five conserved T7SS membrane components, *i.e.* EccB, EccC, EccD and EccE (Ecc stands for <u>E</u>SX <u>c</u>onserved <u>c</u>omponent) and Mycosin or MycP. EccB is a single transmembrane domain (TMD) protein with a large periplasmic domain. EccC is a P-loop NTPase with four FtsK/SpoIIIE-like nucleotide binding domains (NBDs) and the motor protein of the complex, which has been shown to bind substrates (Champion *et al*, 2006; Rosenberg *et al*, 2015). EccD is the most hydrophobic component with eleven TMDs and EccE is anchored in the membrane via two TMDs. The four Ecc components have been shown to assemble into a large ∼2 MDa machinery (Houben *et al*, 2012b) and several structures of (sub)complexes have been solved by electron microscopy (EM) (Beckham *et al*, 2017; Beckham *et al*, 2021; Famelis *et al*, 2019; Poweleit *et al*, 2019). The fifth membrane component MycP, a subtilisin-like protease with a single C-terminal TMD, interacts with and stabilizes the EccBCDE membrane complex (van Winden *et al*, 2020; van Winden *et al*, 2016). The recently solved cryo-EM structure of a full ESX-5 inner membrane complex of *M. tuberculosis* revealed the position of MycP within this complex, confirming the stabilizing role of MycP within the membrane assembly (Bunduc *et al*, 2021).

Most substrates that are secreted by mycobacterial T7SSs belong to three substrate families, *i.e.* the Esx, PE and PPE protein families. In pathogenic mycobacteria, especially the number of *pe* and *ppe* genes has expanded extensively, covering nearly 10% of the coding capacity (Cole *et al*, 1998). A specific feature of T7SS substrates is that they are secreted as folded heterodimers (Ates, 2020). Two different Esx proteins assemble into a secreted heterodimer, whereas PE proteins interact with specific PPE proteins to form a structurally similar substrate pair. Dimerization is mediated by hydrophobic interactions resulting in a conserved four helix bundle structure. The C-terminal flexible domain of one of the secretion partner proteins additionally contains a YxxxD/E secretion signal that is important for secretion (Daleke *et al*, 2012).

Mycobacterial T7SSs also have two cytosolic components, EccA and EspG. EspG is a PPE-dedicated chaperone that binds to a hydrophobic patch on the so-called helical tip domain of these proteins (Ekiert & Cox, 2014; Korotkova *et al*, 2014). Binding of EspG to PPE substrates is required to keep PE/PPE pairs soluble in the cytosol (Ekiert & Cox, 2014; Korotkova *et al*., 2014). EspG is additionally involved in determining system-specific substrate recognition, as swapping the EspG-binding domain between two PPE substrates resulted in rerouting of the ESX-1 substrate PPE68_1 to the ESX-5 system in *Mycobacterium marinum* (Phan *et al*, 2017). EccA is an AAA+ ATPase that has been hypothesized to disrupt the EspG/PPE interaction, leading to recycling of EspG in the cytosol, although the importance of this ATPase for the secretion process remains controversial and could be dependent on growth conditions (Ekiert & Cox, 2014; Phan *et al*, 2018).

Studying the functioning of T7SSs is impaired by the slow growth rate and virulence of pathogenic mycobacteria. Certain systems, such as ESX-3 and ESX-5, are also essential for growth, complicating genetic analyses (Ates *et al*., 2015; Serafini *et al*, 2009; Siegrist *et al*., 2009). Furthermore, studying the potential role of specific substrates in the secretion process is hampered by the redundancy between highly similar substrates (Ates, 2020; Gey van Pittius *et al*, 2006). *Mycobacterium smegmatis*, an avirulent and fast-growing mycobacterium, is commonly used as a model for studying mycobacterial physiology. This species possesses ESX-1, ESX-3 and ESX-4, but lacks the ESX-2 and ESX-5 systems, which are exclusively found in slow-growing mycobacteria.

Here, we exploited an ESX-5 system of *Mycobacterium xenopi* that we have previously functionally reconstituted in *M. smegmatis* (Beckham et al., 2017) to study the role of individual *esx-5* genes in system functionality. We took advantage of the limited number of substrate genes present in the introduced *esx-5* locus and defined the minimal number of genes necessary for successful assembly of the ESX-5 membrane complex and ESX-5 mediated secretion.

## Results

### The inter-dependence of ESX-5 components for stable expression in *M. smegmatis*

To be able to efficiently dissect the individual roles of T7SS components, we used the ESX-5 system of the slow-growing, moderately thermophilic and pathogenic mycobacterial species *M. xenopi* (Fig. 1A), which we have previously functionally reconstituted in *M. smegmatis* (Beckham *et al*., 2017; van Winden *et al*., 2020). As *M. smegmatis* does not have and never had a native ESX-5 system, the only ESX-5 substrates are those encoded by the introduced *esx-5* locus, *i.e.* the Esx pair EsxM/EsxN and two PE/PPE pairs that are specific for *M. xenopi* (Fig. 1A). This is in contrast to ESX-5 systems in natural hosts, which secrete multiple Esx pairs and a large number of PE and PPE substrates, of which some show redundancy (Abdallah *et al*, 2007; Ates, 2020). The limited number of substrates secreted by the reconstituted system provides an excellent opportunity to analyze their individual roles in T7SS functioning, bypassing any potential redundancy issues.

**Figure 1.**
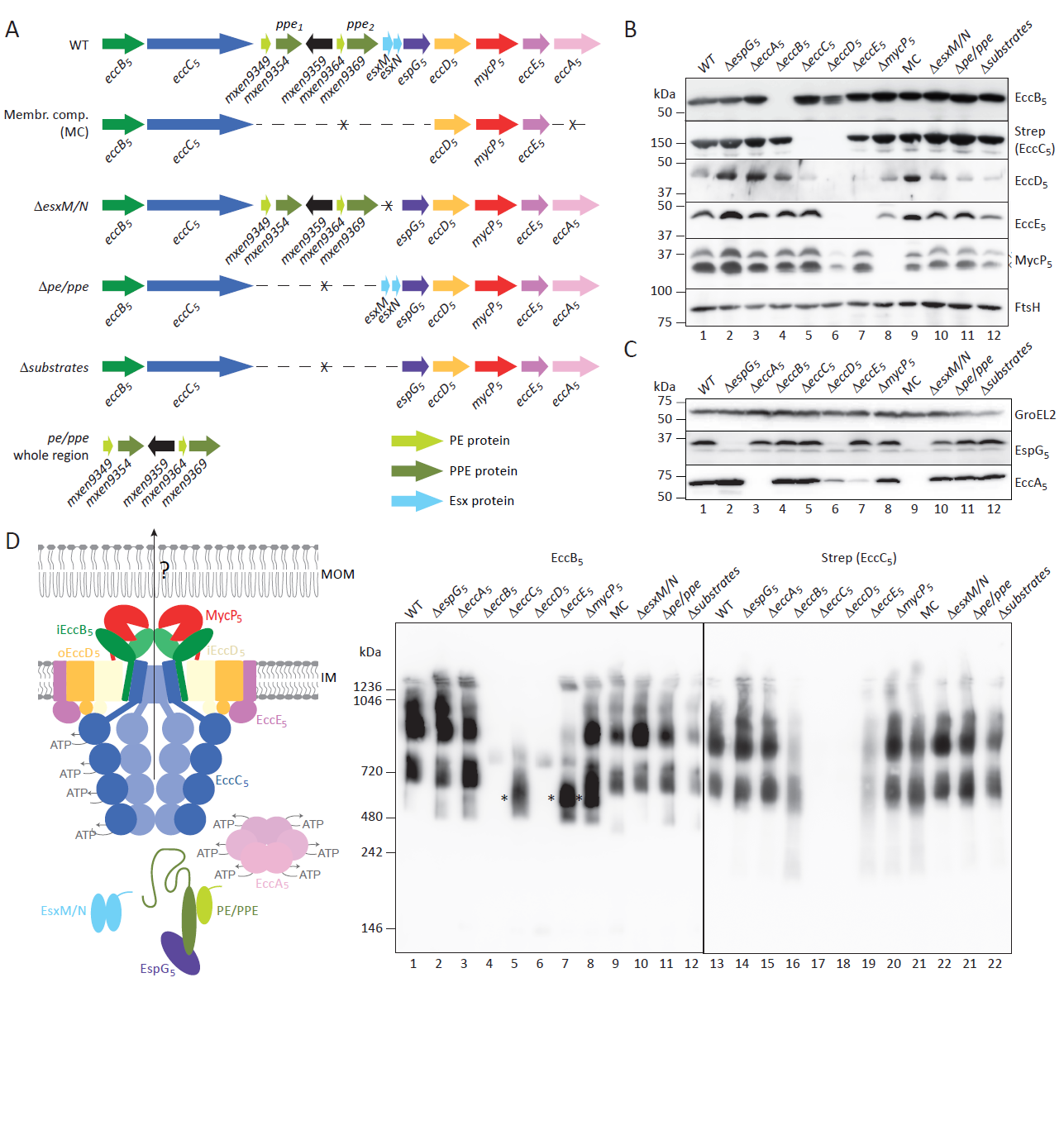
Protein expression and membrane complex formation for ESX-5 mutants. (A) Genetic organization of the *esx-5* locus of *M. xenopi* RIVM700367 and several of its derivatives. Depicted are the complete gene cluster, the cluster derivatives, in which more than one gene was deleted, and the gene organization for the *pe*/*ppe* region complementation plasmid. (B) SDS-PAGE and immunoblot analysis of cell envelope fractions of *M. smegmatis* carrying the ESX-5_Mxe_ plasmid or its derivatives using antibodies reactive against the four membrane components EccB_5_, EccD_5_, EccE_5_ and MycP_5_ or antibodies directed against a Strep-tag linked to the membrane component EccC_5_. Antibodies against the unrelated membrane protein FtsH were used as loading control. Notably, MycP_5_ is processed in one of the loops of its protease domain (van Winden *et al*., 2016), resulting in two immuno-reactive bands (arrow heads). The high hydrophobicity of EccD_5_ compromises its biochemical analysis, resulting in its variable detection by SDS-PAGE and immunoblotting. (C) SDS-PAGE and immunoblot analysis of whole cell lysates of *M. smegmatis* carrying the ESX-5_Mxe_ plasmid or its derivatives using antibodies reactive against the cytosolic ESX-5 components EccA_5_ and EspG_5_. Antibodies against GroEL2 were used as loading control. (D) Model of the ESX-5 system depicting all components and substrates (left) and BN-PAGE and immunoblot analysis of DDM-solubilized cell envelope fractions of *M. smegmatis* carrying the ESX-5_Mxe_ plasmid or its derivatives using antibodies against EccB_5_ or the Strep-tag introduced on EccC_5_ (right). Asterisks denote the ∼500 kDa subcomplex seen in the *eccC_5_, eccE_5_* and *mycP_5_* mutants. MOM, mycobacterial outer membrane; IM, inner membrane.

We exploited this non-essential reconstituted system to make single deletions of every *esx-5* gene that encodes for system components, *i.e.* the *eccABCDE*, *mycP* and *espG* genes. In addition, we created *esx-5* plasmids that were devoid of all *pe* and *ppe* genes (*Δpe/ppe*), lacking the two *esx* genes (*ΔesxM/N*) or devoid of all genes that encode for substrates (*Δsubstrates*) (Fig. 1A). The *Δsubstrates* and *Δpe/ppe* plasmids included the deletion of a non-conserved gene situated between the *pe/ppe* pairs, *i.e. mxen9359,* which codes for a putative SAM-methyltransferase (Fig. 1A). Finally, we also created a construct that only contained the five conserved membrane components (MC) (Fig 1A). These constructs were used to assess the effect of each gene deletion on the protein levels of other components, on the formation of the ESX-5 membrane complex and on ESX-5 mediated secretion.

First, we analyzed the impact of the various gene deletions on the expression of the other system components by SDS-PAGE and immunoblotting, using specific antisera for the different ESX-5 components. We used isolated cell envelope fractions of *M. smegmatis* bearing the different *esx-5* plasmids to detect membrane components (Fig. 1B) and whole cell lysates of the same strains to assess protein levels of the cytosolic components (Fig. 1C). Deletion of the genes coding for the cytosolic components EspG_5_ and EccA_5_ did not affect the protein levels of any of the five inner membrane components, *i.e.* EccB_5_, EccC_5_ (strep tagged), EccD_5_, EccE_5_ and MycP_5_ (Fig. 1B). Furthermore, in the absence of EccA_5_, the protein levels of EspG_5_ were not affected and *vice versa* (Fig. 1C). Deleting the membrane components EccB_5_, EccC_5_ or MycP_5_ also did not affect protein levels of the four other membrane components and the two cytosolic components. In contrast, the absence of EccE_5_ seems to affect protein levels of EccD_5_ and also of EccA_5_, although the latter could be attributed to polar effects of the gene deletion, as *eccA_5_* lies downstream of *eccE_5_*. Knocking out *eccD_5_* showed the most severe effect on the protein levels of the other components. While EccC_5_ and EccE_5_ were undetectable by immunoblotting in the absence of EccD_5_, the protein levels of the membrane components EccB_5_ and MycP_5_ were reduced. Strikingly, the two cytosolic components were also affected by the *eccD_5_* deletion, as EspG_5_ could not be detected anymore and levels of EccA_5_ were severely reduced. In conclusion, deletion of the cytosolic components does not impact the protein levels of any of the other components, while deleting *eccD_5_* severely impacted the levels of all components.

### Only the five membrane components are required for full assembly of the ESX-5 membrane complex

Next, we investigated which components play a role in the successful assembly of the ESX-5 inner membrane complex. For this, we analyzed DDM-solubilized cell envelope fractions by blue native PAGE (BN-PAGE), followed by immunoblotting using our anti-EccB_5_ antibody or an antibody that recognized the Strep-tag on EccC_5_ (Fig 1D). Similar to previous observations (Beckham *et al*., 2017), BN-PAGE analysis of samples from *M. smegmatis* bearing the complete *esx-5* locus resulted in three main complexes that reacted with the anti-EccB_5_ antibody, the largest complex representing the full complex of ∼2 MDa and two subcomplexes of ∼900 kDa and ∼700 kDa. Staining with the anti-Strep antibody showed a similar immunoblot pattern. Taking into consideration the influence of the detergent, lipids and the Coomassie dye on the size estimation of membrane proteins by BN-PAGE (Wittig *et al*, 2010), these three complexes could represent the full hexameric complex, the dimeric subcomplex and an individual protomer, respectively. In the absence of the cytosolic components EccA_5_ and EspG_5_, membrane complex assembly was not affected, as the BN-PAGE immunoblot pattern in their absence was similar as for the WT construct (Fig. 1D). Similarly, none of the substrate deletion plasmids showed differences in ESX-5 complexes, suggesting that substrates do not play a role in the assembly and/or stability of the ESX-5 membrane complex. In line with this observation, a plasmid containing only the five *esx-5* genes that code for the inner membrane components resulted in membrane complex formation similar to the intact *esx-5* locus.

Deletion of the five membrane component genes, *i.e. eccB_5_*, *eccC_5_*, *eccD_5_*, *eccE_5_* and *mycP_5_*, showed varying effects on complex formation. In line with the observation that the *eccD_5_* deletion severely affected expression of the four other membrane components, no complex formation could be observed in this condition. This suggests that EccD_5_ is crucial for complex formation and/or stability, which is in line with its central location within the protomers of the membrane complex (Fig. 1D). While deletion of either *eccB_5_*, *eccC_5_* or *eccE_5_* did not drastically affect the expression of the other membrane components (Fig. 1B), the assembly of the membrane complex, as seen by BN-PAGE analysis, was affected. Deletion of *eccB_5_* affected formation of especially the full ∼2 MDa complex, which is in line with previous findings that EccB_5_ is involved in complex multimerization (Bunduc *et al*., 2021; Famelis *et al*., 2019). In the absence of the central ATPase EccC_5_, which gates the potential secretion pore with its TMDs, membrane complex formation was almost completely abolished, with this construct exhibiting only a ∼500 kDa subcomplex containing EccB_5_, which was not observed with the WT construct. Deleting *eccE_5_* showed a reduced amount of complexes, using both antibodies, which is in line with our previous observation that a lower number of full complexes could be isolated under this condition (Beckham *et al*., 2017). In addition, the ∼500 kDa complex, also present with the Δ*eccC_5_* construct, was observed in the absence of EccE_5_ using the anti-EccB_5_ but not the anti-Strep antibody. Finally, while the *ΔmycP_5_* plasmid showed no major differences in the formation of the inner membrane complex, the ∼500 kDa EccB_5_-containing complex was also observed here. These results were similar to previous observations in *M. marinum*, where MycP_1_ and MycP_5_ of the ESX-1 and ESX-5 system, respectively, have been reported to be involved in stabilizing its corresponding T7SS membrane complex (van Winden et al., 2016).

Taken together, optimal assembly of the ESX-5 membrane complex requires only the five membrane components.

### *esx-5* locus-encoded PE/PPE proteins are required for secretion of Esx proteins

We previously observed that introduction of the plasmid that contains the complete *esx-5* locus of *M. xenopi* in *M. smegmatis* resulted in the secretion of the locus-encoded ESX-5 substrate EsxN and also of *M. marinum* PPE18, which was introduced by an additional plasmid containing the gene pair *pe31/ppe18* (Beckham *et al*., 2017). This shows that the reconstituted system is functional. We therefore tested next the impact of the various gene deletions on the ability of this system to secrete EsxN and PPE18, the latter by again introducing the plasmid containing the *pe31/ppe18* of *M. marinum* (PPE18 containing a C-terminal HA-tag for detection) (Fig. 2A).

**Figure 2.**
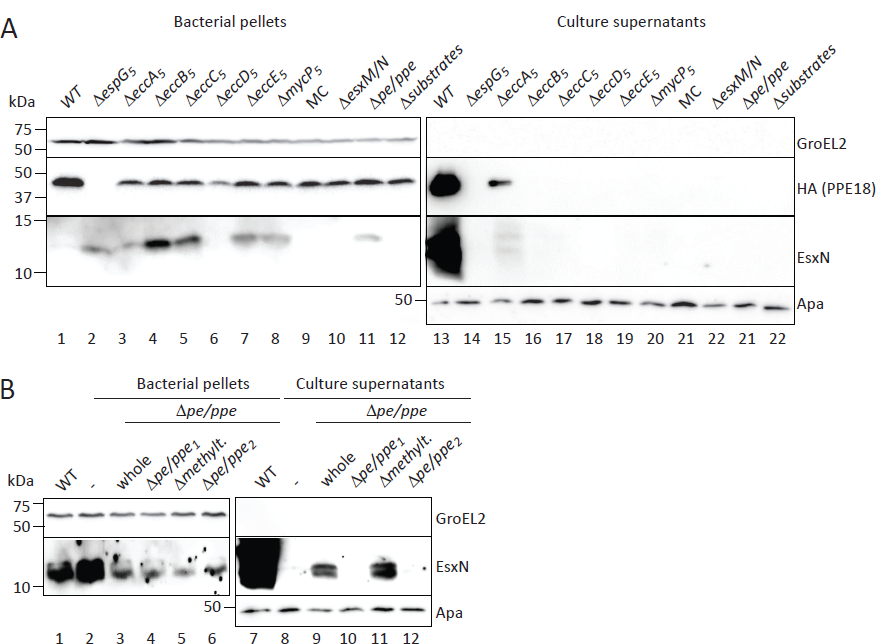
Secretion analysis of *M. smegmatis* carrying the ESX-5_Mxe_ plasmid or its derivatives. (A) SDS-PAGE and immunoblot analysis of secreted fractions (culture supernatants) and whole cell lysates (bacterial pellets) using antibodies against EsxN. In addition, an HA-antibody was used to detect PPE18-HA that was encoded together with PE31, its partner protein, on a separate plasmid. Antibodies against GroEL2 were used as lysis and loading control for the whole cell lysates and antibodies against Apa were used as loading control for the secreted fraction. (B) Secretion analysis of *M. smegmatis* carrying the ESX-5_Mxe_ Δ*pe/ppe* plasmid and various *pe/ppe* complementation vectors. The used bacterial fractions and antibodies are the same as under A.

While both EsxN and PPE18-HA were efficiently secreted with the WT construct, they were also secreted in the absence of *eccA_5_*, albeit at a reduced level (Fig. 2A). Deletion of the other cytosolic component *espG_5_* blocked the secretion of both PPE18-HA and EsxN (Fig. 2A). This critical role of EspG_5_ for successful secretion of both the PE/PPE proteins and EsxM/EsxN is in line with previous findings (Korotkova *et al*., 2014). Additionally, PPE18-HA was not present in the bacterial pellet, suggesting that EspG_5_ is essential for the stability of this protein, as observed before for this and other PPE proteins (Ekiert & Cox, 2014; Korotkova *et al*., 2014; Phan *et al*., 2017). In contrast, in the presence of only the membrane components, which did not show secretion of PPE18-HA, this substrate could be detected in the pellet fraction. The instability of PPE18 in the absence of EspG_5_ can be rescued by deletion of *eccA_5_* and/or the substrate-encoded *esx-5* genes. As all five membrane component deletions showed a disruption in membrane complex assembly, it is no surprise that they also abolished the secretion of both EsxN and PPE18-HA. Notably, deleting *eccD_5_* negatively affected also the protein levels of not only PPE18-HA, which is in line with the absence of EspG_5_ under this condition (Fig. 1C), but also of EsxN. In the absence of EsxM/EsxN, secretion but not the stable expression of PPE18-HA was abrogated, showing that this Esx heterodimer is required for secretion of the PE/PPE pair (Fig. 3A). Strikingly, in the absence of PE/PPE proteins encoded by the *esx-5* locus, which did not affect expression of any ESX-5 components nor assembly of the ESX-5 membrane complex, secretion but not stable expression of both EsxN and PPE18 was abolished (Fig. 2A). From this, we conclude that the PE/PPE pairs of the *esx-5* locus are important for the secretion of the Esx substrates and at least one other PPE protein. This is in line with a recent observation for ESX-1 in *M. marinum*, where secretion of EsxA is dependent on the co-secretion of PPE68 encoded by the same gene cluster (Cronin *et al*, 2022; Damen *et al*, 2022).

**Figure 3.**
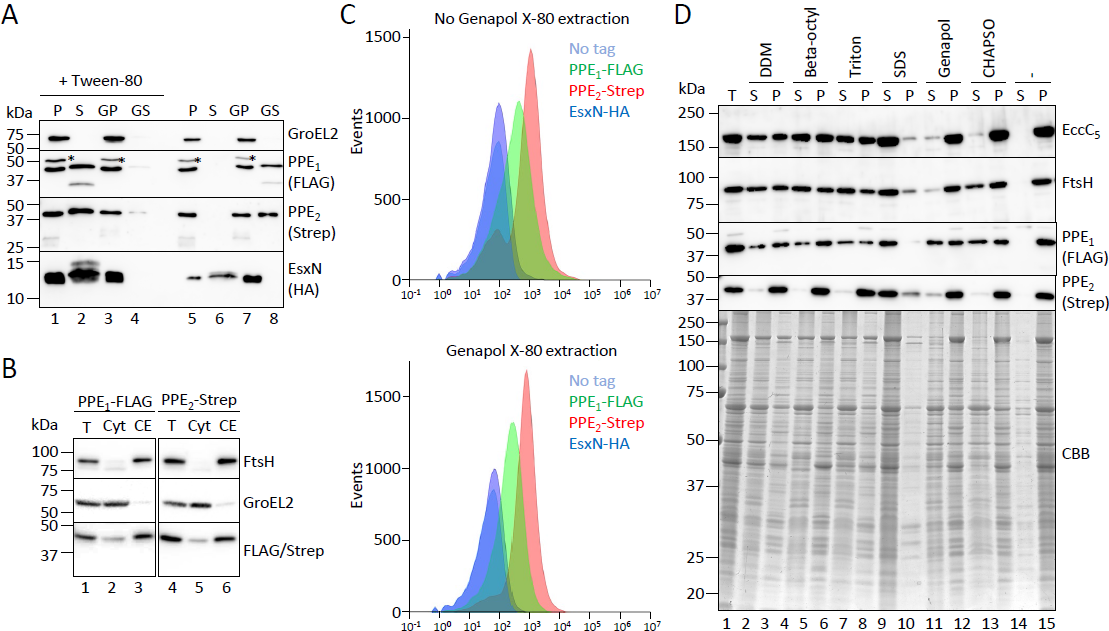
Subcellular localization of the tagged PPE substrates of the *esx-5_Mxe_* locus expressed in *M. smegmatis*. (A) Secretion analysis of *M. smegmatis* carrying the ESX-5_Mxe_ plasmid expressing tagged substrates. SDS-PAGE and immunoblot analysis of secreted fractions (culture supernatants) and whole cell lysates (bacterial pellets) of *M. smegmatis* carrying the ESX-5_Mxe_ plasmid encoding the first PPE substrate (PPE_1_) with a FLAG-tag at the N-terminus, the second PPE protein (PPE_2_) with a Strep-tag at the C-terminus and EsxN with an HA-tag at the C-terminus. Antibodies against GroEL2 were used as lysis control. Cells were grown in the presence or absence of Tween-80, after which they were harvested and half was treated with Genapol X-080 to extract surface proteins. P, bacterial pellet; S, culture supernatant; GP, Genapol pellet; GS, Genapol supernatant. Background bands are marked with asterisks. (B) Cells expressing the *M. xenopi esx-5* locus encoding substrates with tags were grown in the presence of 0.05% Tween 80, after which cells were fractionated into total (T), cytosol (Cyt) and cell envelope (CE) fractions. Fractions were loaded in equal amounts. Antibodies against FtsH (inner membrane protein) and GroEL2 (cytosolic protein) were used as fractionation control. (C) Surface exposure of substrates on whole cells was measured by flow cytometry of cells grown in the presence of Tween-80 without (top) or with (bottom) surface protein extraction with Genapol X-080 after culture harvesting. (D) SDS-PAGE and immunoblot or Coomassie brilliant blue (CBB) stain analysis of cell envelope fractions isolated from cells treated with Genapol X-080 and solubilized with the mentioned detergents. Concentrations for the detergents were: 0.25% DDM, 1% n-Octyl-β-D-Glucopyranoside, 2% Triton X-100, 2% SDS, 0.5% Genapol X-080 and 5% CHAPSO. Fractions were loaded in equal amounts. Antibodies against EccB_5_ and FtsH were used as inner membrane controls. S, soluble fraction; P, non-soluble fraction.

To analyze the role of the individual *pe/ppe* gene pairs and of the putative methyltransferase gene that is located between the two *pe/ppe* pairs, we created four complementation vectors containing either the entire deleted region (whole; Fig. 1A), one of the two *pe/ppe* gene pairs with the methyltransferase gene (*Δpe/ppe_1_* and *Δpe/ppe_2_*) or only the two *pe/ppe* pairs without the methyltransferase gene (*Δmethylt.*). EsxN secretion could be rescued by complementation with a plasmid containing the entire deleted region (Fig. 2B), although the observed secretion levels were reduced when compared to the levels with the WT construct. This reduction could be due to the fact that both introduced plasmids contain the same origin of replication, thereby reducing individual plasmid copy numbers and the overall protein levels. The same (reduced) secretion levels were observed when we complemented with a plasmid that contains the entire deleted region without the gene coding for the predicted methyltransferase, suggesting that this methyltransferase gene is not required for secretion. Methyltransferases are not often associated with *esx-5* or other T7SS loci, suggesting that the presence of this gene could be a coincidence. However, deletion of either of the two *pe/ppe* pairs in the complementation construct abrogated secretion of EsxN completely. This shows that both PE/PPE pairs are required for EsxN secretion. Notably, complementing the *Δpe/ppe* plasmid with the PE31/PPE18-HA-encoding plasmid also did not restore secretion of EsxN (Fig. 2A). This shows that although PPE18-HA is secreted in an ESX-5 dependent manner, it cannot take over the role of the *esx-5* locus-encoded PE/PPE proteins in EsxN secretion, in line with a model in which Esx and PE/PPE proteins are specifically paired for their co-secretion. The observation that both *esx-5*-encoded PE/PPE proteins and EsxM/EsxN are required for PPE18-HA secretion shows that both substrate classes are required for ESX-5 functionality.

### PPE but not Esx substrates localize to the cell envelope fraction

While the interdependency of substrates for secretion can be explained by a previously proposed dual substrate recognition event by the EccC ATPase, which is required for system activation (Bunduc *et al*, 2020b; Rosenberg *et al*., 2015), substrates could additionally be dependent on each other beyond the T7SS inner membrane complex. The dimensions of the core membrane complex dictate that it spans only the inner membrane. How protein translocation takes place beyond the inner membrane, especially across the mycobacterial outer membrane, is unknown, as there is no clear data on the identity of the outer membrane channel or components. As our reconstituted ESX-5 system is functional in *M. smegmatis*, the proteins that drive secretion over this second membrane are likely encoded by the introduced *esx-5* cluster. Within the *esx-5_Mxe_* locus, the proteins that have thus far been regarded as substrates are the only proteins that could mediate translocation through the outer membrane, as all other components are localized either in the inner membrane or in the cytosol. In addition, in recent years it has become clear that various PE/PPE proteins are involved in the uptake of nutrients and localize to the cell envelope, with PPE51 as the main example (Korycka-Machala *et al*, 2020; Wang *et al*, 2020). To address whether the PPE substrates encoded by the *M. xenopi esx-5* locus could be involved in outer membrane transport of proteins, we tagged the first and the second PPE protein with a FLAG- and a Strep-tag, respectively, either at the N- or C-terminus. Additionally, to be able to detect EsxN more efficiently, we added a C-terminal HA-tag to this substrate, which has previously been shown not to interfere with the secretion of its ESX-1 equivalent EsxA (Damen *et al*, 2020). Subcellular fractionation and secretion analysis showed that a FLAG-tag at the C-terminus of the first PPE protein, hereafter called PPE_1_, partially affected secretion of EsxN, whereas placing the tag at the N-terminus had no noticeable effect on EsxN secretion (Fig. EV 1A). The second PPE protein, hereafter called PPE_2_, could not be detected when tagged with a Strep-tag at the N-terminus and using a Strep antibody, while placing the same tag to the C-terminus showed sufficient protein levels (Fig. EV 1A). With both constructs, EsxN was efficiently secreted to the culture supernatant (Fig. EV 1A), suggesting that PPE_2_ with the N-terminal tag is expressed, but that the tag is cleaved off. Finally, we could also efficiently detect secreted EsxN-HA using the HA antibody.

With this information in hand, we made a final construct in which we removed the previously introduced Strep-tag at the C-terminus of EccC_5_ and modified the substrate genes, so that they encode for PPE_1_ with a FLAG-tag at the N-terminus, PPE_2_ with a Strep-tag at the C-terminus and an HA-tag at the C-terminus of EsxN. We first analyzed protein secretion by *M. smegmatis* carrying this construct when grown in the presence or absence of Tween-80, a mild detergent used as anti-clumping agent for mycobacteria grown in liquid culture (Fig. 3A). EsxN was present in the culture supernatant irrespective of the presence of Tween-80. In contrast, both PPE proteins were found in the culture supernatant only when cells were grown in the presence of the detergent. In the absence of Tween-80, these proteins were found on the cell surface instead, as they could be extracted from the cell surface using the mild detergent Genapol X-080 (Fig 3A). This suggests that the presence of Tween-80 triggers the release of these proteins from the cell surface into the culture medium. Notably, when the gene encoding for the central ATPase EccC_5_ was deleted, neither PPE proteins were secreted anymore, showing that their secretion is dependent on a functional ESX-5 system (Fig. EV 2).

Even though we could extract the PPE proteins, a substantial amount of both PPE proteins remained cell-associated after growth with Tween-80 and/or after Genapol X-080 treatment. To investigate the subcellular location of these subpopulations, *M. smegmatis* cells grown in the presence of Tween-80 were lysed and fractionated into a soluble (cytosolic) and insoluble (cell envelope) fraction (Fig. 3B). Intriguingly, both PPE proteins localized mainly to the cell envelope fraction. Furthermore, flow cytometry analysis using anti-FLAG, -Strep and -HA antibodies revealed that both PPE proteins could be efficiently detected on the cell surface of cells grown with Tween-80 (Fig. 3C, Fig. EV 1B). In contrast, EsxN could not be detected via its C-terminal HA-tag, which is in line with the observation that this substrate is mainly secreted into the culture medium. Interestingly, after treatment of the cells with Genapol X-080, similar levels of surface labeling could be observed as compared to cells that were not extracted with Genapol X-080 (Fig. 3C). This suggests that a significant portion of the cell-associated population of the two PPE proteins is firmly attached to the cell surface. To characterize the cell envelope association of these two PPE proteins further, we subjected isolated cell envelope fractions from Genapol X-080-treated cells to solubilization with a range of detergents, some of which have been used previously for the characterization of mycobacterial membrane proteins (Hoffmann *et al*, 2008; Målen *et al*, 2010). While our inner membrane protein controls EccC_5_ and FtsH could be extracted by the nonionic detergents DDM, beta-octylglucoside and Triton X-100, and the anionic detergent SDS, but not by Genapol X-080 or the zwitter ionic detergent CHAPSO, PPE_1_ could be solubilized, partially or fully, by all these detergents (Fig. 3C). In contrast, PPE_2_ could not be solubilized by any of these detergents, with the exception of SDS. Together, we conclude that the majority of the two *esx-5* locus-encoded PPE proteins of *M. xenopi* localizes to the cell envelope of *M. smegmatis*, where they are surface exposed. While a subpopulation of the PPE proteins associates with the cell surface in a Tween-80/Genapol X-080 sensitive manner, a substantial amount is more firmly attached to the cell surface. The observation that the second PPE protein cannot be extracted from the cell envelope with detergents that are able to solubilize inner membrane proteins, a characteristic that is also observed for known or predicted integral outer membrane proteins (Heinz & Niederweis, 2000; Speer *et al*, 2015), indicates that this protein is integrated into the mycobacterial outer membrane. This protein could therefore potentially form a channel that mediates protein transport across the second membrane of the mycobacterial cell envelope.

## Discussion

By using a functionally reconstituted ESX-5 system in *M. smegmatis*, which naturally lacks this system, we were able to study the role of each *esx-5* gene in component stability, assembly of the membrane embedded machinery and system functionality. The obtained results are summarized in Table 1.

**Table 1.** Overview of results.

| Construct | Component/substrate protein levels |  |  |  | Complex formation | Secretion |  |
| --- | --- | --- | --- | --- | --- | --- | --- |
|  | Membrane | Cytosolic | EsxN | PPE18 |  | EsxN | PPE18 |
| WT | ++ | ++ | ++ | ++ | ++ | ++ | ++ |
| $\Delta espG_5$ | ++ | ++ | + | - | ++ | - | - |
| $\Delta eccA_5$ | ++ | ++ | + | + | ++ | + | + |
| $\Delta eccB_5$ | ++ | ++ | + | + | - | - | - |
| $\Delta eccC_5$ | + | ++ | + | + | - | - | - |
| $\Delta eccD_5$ | - | +/- | - | +/- | - | - | - |
| $\Delta eccE_5$ | + | +/- | + | + | + | - | - |
| $\Delta mycP_5$ | ++ | ++ | + | + | + | - | - |
| MC | ++ | n.a. | n.a. | + | ++ | n.a. | - |
| $\Delta esxM/N$ | ++ | ++ | n.a. | + | ++ | n.a. | - |
| $\Delta pe/ppe$ | ++ | ++ | + | + | ++ | - | - |
| $\Delta substrates$ | ++ | ++ | n.a. | + | ++ | n.a. | - |

-, entirely affected; +/-, significantly affected; +, slightly affected; ++, not affected; n.a, not applicable

Deletion of individual membrane components showed varying degrees of defects in membrane complex assembly, while functionality of the ESX-5 system in all these mutants was completely abolished. An *eccD_5_* mutant showed the most striking phenotype. Here, membrane complex assembly was completely abolished, highlighting its role as a scaffold for the assembly of the inner membrane complex. This mutant also exhibited highly affected protein levels, not only for the remaining membrane components but also for cytosolic chaperones and substrates. The fact that the *esx-5* gene cluster is introduced in *M. smegmatis* that lacks ESX-5 itself makes it unlikely that these reduced levels are caused on a transcriptional level by a negative feedback loop, as has been described for the ESX-1 system (Abdallah *et al*, 2019; Bosserman *et al*, 2017; Chen *et al*, 2016). It suggests that the absence of EccD_5_ triggers a domino-like effect on the stability of the other proteins, starting with an unassembled inner membrane machinery, which leads to degradation of the remaining membrane components and, in the absence of the membrane complex, also of the components found in the cytosol, hereby destabilizing the substrates.

Neither substrate nor chaperone mutants showed a defect in assembly of the membrane complex and complex assembly required only the five conserved membrane components. Nevertheless, conformational changes which cannot be accounted for by the methods used here could still occur. While functionality was tightly connected with the successful assembly of the membrane complex, the minimal functional T7SS ESX-5 unit requires the five membrane components, EspG_5_, both PE/PPE pairs that are encoded by the *esx-5* locus and an Esx heterodimer. EspG_5_ proved to be critical for successful secretion of both PE/PPE and EsxM/N proteins, similar to previous findings (Korotkova *et al*., 2014). The system without EccA_5_ was still functional, although to a lower extent. EccA proteins have been shown to be required for secretion in some but not all studies (Bottai *et al*, 2012; Joshi *et al*, 2012; Phan *et al*., 2018).

In addition, we showed that secretion of the Esx heterodimers by ESX-5 is dependent on the locus encoded PE/PPE proteins. This is in line with a recent observation for ESX-1 in *Mycobacterium marinum*, where secretion of EsxA is dependent on the co-secretion of PPE68 that is encoded by the same gene cluster(Cronin *et al*., 2022; Damen *et al*., 2022). Notably, PPE68 was observed to be degraded on the cell surface upon export. Since PE31/PPE18 was not able to complement secretion of the Esx pair, our results are in line with a model in which Esx and PE/PPE proteins are specifically paired for their co-secretion. The observation that both *esx-5* encoded PE/PPE proteins and EsxM/EsxN are required for PPE18-HA secretion shows that both substrate classes are necessary for ESX-5 functionality.

Substrate interdependence for secretion, which has been observed also in other T7SSs (Cronin *et al*., 2022; Damen *et al*., 2022), can be explained by previously proposed models in which the membrane complex exhibits two separate recognition events, one between NBD3 of EccC and EsxB and one mediated by the linker2 domain of EccC with PE/PPE proteins (Bunduc *et al*., 2020b; Rosenberg *et al*., 2015). Substrate interdependency for secretion can also be a phenomenon taking place beyond the inner membrane. As secretion of all three heterodimeric substrates across the full cell envelope is dependent on the reconstituted ESX-5 system in our *M. smegmatis* model organism, all components required for secretion across the cell envelope, including the outer membrane, are likely encoded on the *esx-5_xen_* plasmid. Excluding inner membrane components and cytosolic chaperones leaves only the substrates to be responsible for driving secretion over the outer membrane. Indeed, both PPE substrates, but not EsxN, localize to the cell envelope fraction and are surface exposed, implying they are embedded in the outer membrane. Furthermore, especially PPE_2_, showed a phenotype consistent with known mycobacterial outer membrane proteins, in that it cannot be readily solubilized by detergents. Combined with their essential role in secretion of the Esx heterodimer and the exogenous PE31/PPE18 pair, it suggests a new role for these specific PPE proteins in the transport of proteins across the mycolic acid containing layer.

Based on the combined results, we propose two working models for the role of these *esx-5-* encoded PPE proteins. In the first model, the roles of the two PPE proteins in secretion are different, considering the differences in membrane association. Here, the first PPE protein could play a role in activation of the inner membrane complex and co-secretion together with the EsxM/EsxN heterodimer, similar as has been proposed for the ESX-1 dependent Esx and PE/PPE substrates (Cronin *et al*., 2022; Damen *et al*., 2022), while the second PPE protein could form the channel for protein translocation over the outer membrane. In the second model, the two PPE proteins form the outer membrane channel together, with PPE_2_ creating the integral part and PPE_1_ forming a peripheral portion of the complex. As these PPE proteins are specific for *M. xenopi*, further studies are required to determine whether the *esx-5*-associated PPE proteins of other mycobacterial species, such as *M. tuberculosis*, have similar properties.

## Materials and methods

### Bacterial strains and growth conditions

*E. coli* Dh5α was grown in LB media supplemented with appropriate antibiotics at 37°C and 200 RPM. *M. smegmatis* MC^2^155 was grown in LB media supplemented with 0.5% Tween 80 (Merck) and appropriate antibiotics at 37°C and 90 RPM. Antibiotics were used with the following concentrations: hygromycin 50 mg/l, streptomycin 30 mg/ml.

### Molecular cloning

Cloning was performed using *E. coli* Dh5α and restriction enzymes from New English Biolabs. Polymerase chain reaction (PCR) products were amplified with IProof DNA polymerase from BioRad and fused in multi-product reactions with InFusion HD from TakaraBio. For construction of plasmid derivatives, genes were deleted, either completely if they were in a single operon or ∼100-150 bp were left at the C-terminus, if these were in an operon with other down- and/or upstream genes. Generally, genes were deleted using the WT ESX-5_Mxe_ plasmid as template using the closest unique restriction sites up- and downstream of the gene of interest and amplification of two or more PCR fragments with compatible 18 bp ends, where the gene of interest was omitted. Because for different deletions the same restriction sites were used, some primers were used for construction of multiple derivatives. For a complete overview of restriction enzymes, primers, plasmids and cloning strategies see Table EV 1, Table EV 2 and Table EV 3.

### Protein secretion and immunoblot analysis

*M. smegmatis* MC^2^155 with plasmids coding for ESX-5_Mxe_ WT or its derivatives were grown in LB liquid medium with 0.05% Tween 80 and appropriate antibiotics at 37°C and 90 RPM until mid-log phase. Cells were sub-cultured at an OD_600_ of 0.05 in LB liquid media under the same conditions for 12-16 hours until an OD_600_ of 0.8-1 was reached. Cultures were spun down, supernatants were passed through 0.2 µm filters and precipitated with trichloroacetic acid (TCA), resulting in culture supernatant fractions. Cell pellets were broken via bead-beating with glass beads, resulting in whole cell lysates or bacterial pellet fractions. SDS loading buffer was added to all samples, boiled and loaded on SDS-PAGE gels (10% or 16% depending on the protein size), transferred to nitrocellulose membranes and stained with appropriate antibodies. Antibodies against EccA_5_, EccB_5_, EccD_5_, EccE_5_, EspG_5_, FtsH, MycP_5_, EsxN (Mtb9.9) and GroEL2 have been described previously (Houben *et al*., 2012b); HA-, FLAG and Strep-antibodies were purchased from Thermo Scientific (2-2.2.14), Sigma-Aldrich (F-1804) and Novusbio (NBP2-41073).

### Cell envelope isolation

*M smegmatis* MC^2^155 with the various ESX-5_Mxe_ plasmids were grown as above to an OD_600_ of ∼1.5. Cells were washed in PBS, resuspended in buffer (50 mM Tris-HCl, 300 mM NaCl and 10% glycerol) and broken by passing two times through a high-pressure homogenizer at 0.3 kbar (Stansted). Unbroken cells were pelleted at 5000xg. Cell envelopes were separated from the soluble fraction by ultracentrifugation at 150.000xg. After ultracentrifugation, pellets were washed and resuspended in buffer and snap-frozen in liquid nitrogen. Where stated, after culture harvesting, cells were incubated with mixing at room temperature with 0.5% Genapol X-80 for 30 minutes, to remove cell surface proteins of bacteria. Subsequently, cells were centrifuged at 5000xg, washed one time with buffer and fractionated as explained above.

### BN-PAGE

For BN-PAGE analysis of membrane complexes, cell envelopes were solubilized with 0.25% DDM for one hour at 4°C. Non-solubilized material was pelleted by centrifugation at 100.000xg for 20 minutes. The resulting supernatant samples were run on 3-12% NativePage Bis-Tris Protein Gels (Invitrogen) with addition of NativePage 5% G-250 Sample Additive (Invitrogen). Gels were blotted to PVDF membrane and stained with appropriate antibodies.

### Flow cytometry analysis

*M. smegmatis* expressing different plasmids was grown to an OD600 of 0.8 to 1.2 in LB with or without Tween-80. Bacteria were pelleted, washed with PBS with 1% bovine serum albumin (BSA; Sigma), and incubated for 2h with antibodies recognizing the FLAG-tag (F-1804; Sigma-Aldrich), Strep-tag (NBP2-41073; Novusbio) or influenza virus hemagglutinin tag (2-2.2.14; Thermo Scientific). After washing with PBS with 1% BSA, bacteria were incubated with secondary goat anti-mouse IgG (A-21235; ThermoFischer) or secondary goat anti-rabbit IgG (A-21244; ThermoFischer) conjugated to Alexa 647 antibodies for 30 min. After washing with PBS with 1% BSA, bacteria were analyzed by flow cytometry (Attune NxT; ThermoFisher). As a control, bacteria were incubated only with secondary antibodies.

## Acknowledgements

We thank Roy Ummels for technical help and advice. This work received funding by a VIDI grant (864.12.006; to C.M.B. and E.N.G.H.) from the Netherlands Organization of Scientific Research. This project has received funding from the European Union’s Horizon 2020 research and innovation programme under the Marie Sklodowska-Curie grant agreement no. 101030373 (to C.M.B.). This project was supported by funds available to T.C.M. through the Behörde für Wissenschaft, Forschung und Gleichstellung of the city of Hamburg at the Institute of Structural and Systems Biology at the University Medical Center Hamburg–Eppendorf (UKE). The laboratory of T.C.M. is supported by DESY (German Electron Synchrotron Center).

---

**Expanded View Figure 1.**
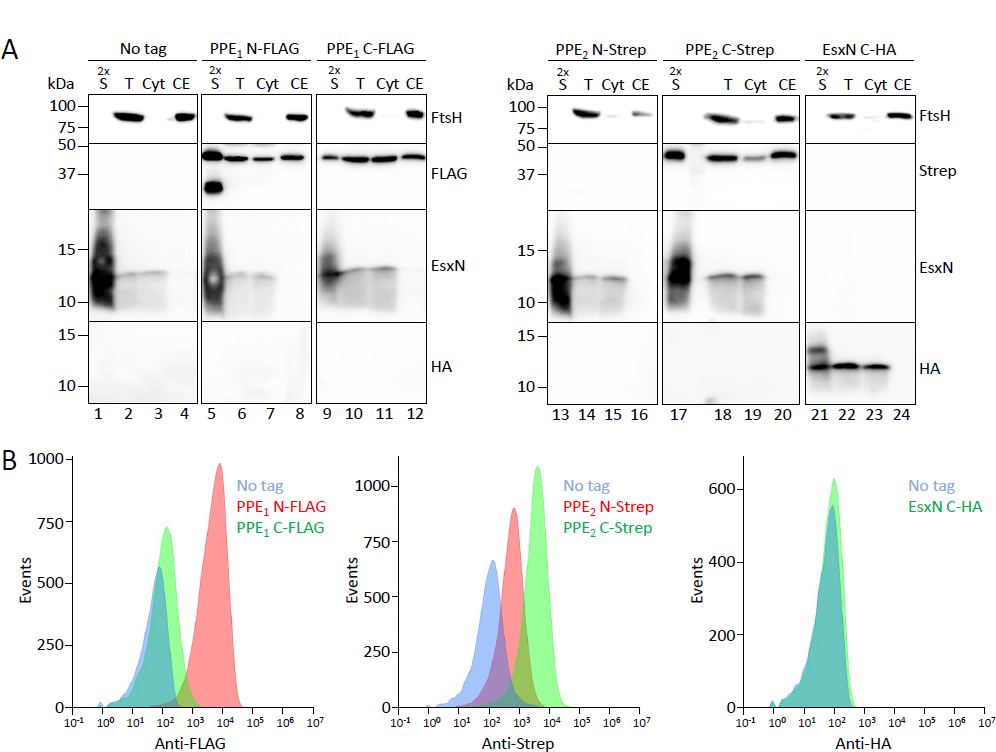
Subcellular fractionation of *M. smegmatis* carrying the *esx-5_Mxe_* plasmid expressing tagged ESX-5 substrates. (A) Cells expressing the *M. xenopi esx-5* locus encoding substrates with tags at the N or C-termini were grown in the presence of 0.05% Tween 80, after which supernatants (S) were separated from bacterial cells. The cells were subsequently fractionated into total (T), cytosol (Cyt) and cell envelope (CE) fractions. Fractions were loaded in a ratio of 2:1:1:1 for S:T:Cyt:CE. Antibody against FtsH (inner membrane protein) was used as fractionation control. (B) Surface exposure of substrates on whole cells was measured by flow cytometry of cells grown in the presence of Tween-80.

**Expanded View Figure 2.**
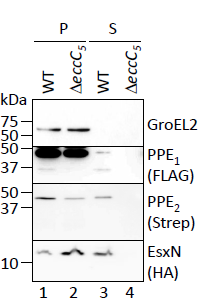
Secretion analysis of the *esx-5-*encoded PPE proteins by *M. smegmatis* carrying the *esx-5_Mxe_* Δ*eccC_5_* plasmid. SDS-PAGE and immunoblot analysis of secreted fractions (S) and whole cell lysates (P) of *M. smegmatis* carrying the ESX-5_Mxe_ plasmid encoding the first PPE substrate (PPE_1_) with a FLAG-tag at the N-terminus, the second PPE protein (PPE_2_) with a Strep-tag at the C-terminus and EsxN with a HA-tag at the C-terminus and the same plasmid with *eccC_5_* deleted. Antibodies against GroEL2 were used as lysis control. Cells were grown in the presence of Tween-80.

**Expanded View Table 1.** List of plasmids used in this study

**Expanded View Table 2.** Primers used in this study for molecular cloning

**Expanded View Table 3.** Overview of molecular cloning strategies for generating the plasmids shown in Table EV1 and using primers shown in Table EV2.

**Expanded View Table 1:** List of plasmids used in this study.

| Nr. | Construct | Source |
| --- | --- | --- |
| 1. | pMV *esx-5*<sub>Mxe</sub> | Beckham *et al.*, 2017 |
| 2. | pMV *esx-5*<sub>Mxe</sub> Δ*eccB*<sub>5</sub> | this study |
| 3. | pMV *esx-5*<sub>Mxe</sub> Δ*eccC*<sub>5</sub> | this study |
| 4. | pMV *esx-5*<sub>Mxe</sub> Δ*pe/ppe* | this study |
| 5. | pMV *esx-5*<sub>Mxe</sub> Δ*esxM/N* | this study |
| 6. | pMV *esx-5*<sub>Mxe</sub> Δ*substrates* | this study |
| 7. | pMV *esx-5*<sub>Mxe</sub> Δ*espG*<sub>5</sub> | this study |
| 8. | pMV *esx-5*<sub>Mxe</sub> Δ*eccD*<sub>5</sub> | this study |
| 9. | pMV *esx-5*<sub>Mxe</sub> Δ*mycP*<sub>5</sub> | van Winden *et al.*, 2020 |
| 10. | pMV *esx-5*<sub>Mxe</sub> Δ*eccE*<sub>5</sub> | this study |
| 11. | pMV *esx-5*<sub>Mxe</sub> Δ*eccA*<sub>5</sub> | this study |
| 12. | pMV *esx-5*<sub>Mxe</sub> membrane components (MC) | this study |
| 13. | pSMT3 *pe/ppe* whole | this study |
| 14. | pSMT3 *pe/ppe* Δ*pe/ppe*<sub>1</sub> | this study |
| 15. | pSMT3 *pe/ppe* Δmethyltransferase | this study |
| 16. | pSMT3 *pe/ppe* Δ*pe/ppe*<sub>2</sub> | this study |
| 17. | pMV *esx-5*<sub>Mxe</sub> *ppe*<sub>1</sub> FLAG N-term | this study |
| 18. | pMV *esx-5*<sub>Mxe</sub> *ppe*<sub>1</sub> FLAG C-term | this study |
| 19. | pMV *esx-5*<sub>Mxe</sub> *ppe*<sub>2</sub> Strep N-term | this study |
| 20. | pMV *esx-5*<sub>Mxe</sub> *ppe*<sub>2</sub> Strep C-term | this study |
| 21. | pMV *esx-5*<sub>Mxe</sub> *esxN-HA* | this study |
| 22. | pMV *esx-5*<sub>Mxe</sub> *ppe*<sub>1</sub> FLAG N-term, *ppe*<sub>2</sub> Strep C-term, *esxN-HA*, No Strep on *eccC*<sub>5</sub> | this study |
| 23. | pMV *esx-5*<sub>Mxe</sub> *ppe*<sub>1</sub> FLAG N-term, *ppe*<sub>2</sub> Strep C-term, *esxN-HA*, Δ*eccC*<sub>5</sub> | this study |

**Expanded View Table 2:**
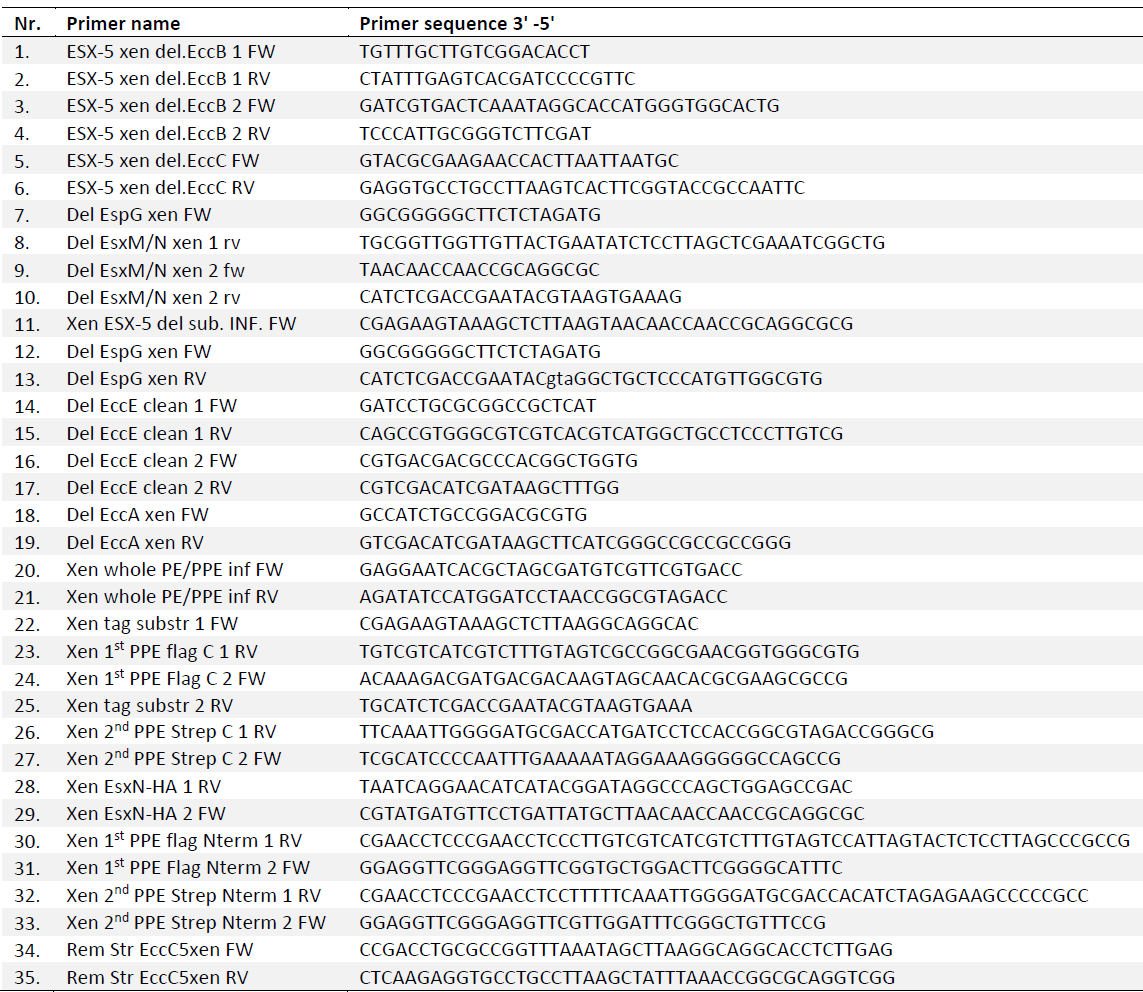
List of primers used in this study.

**Expanded View Table 3:** Overview of molecular cloning strategies for generating the plasmids shown in Expanded View Table 1 and using primers shown in Expanded View Table 2.

| Nr. | Construct | Starting backbone | Restriction sites used to cut the backbone | Primers used | PCR product size | Comments |
| --- | --- | --- | --- | --- | --- | --- |
| 2. | pMV *esx-5<sub>Mxe</sub> ΔeccB<sub>5</sub>* | pMV *esx-5<sub>Mxe</sub>* | EcoRV, SexAI | 1, 2<br>3, 4 | 330 bp<br>1140 bp | Restricted backbone and the two PCR products were ligated with InFusion HD. |
| 3. | pMV *esx-5<sub>Mxe</sub> ΔeccC<sub>5</sub>* | pMV *esx-5<sub>Mxe</sub>* | PacI, AflIII | 5, 6 | 2174 bp | Restricted backbone and the PCR product were ligated with InFusion HD. |
| 4. | pMV *esx-5<sub>Mxe</sub> Δpe/ppe* | pMV *esx-5<sub>Mxe</sub>* | AflIII, XbaI |  |  | Restriction removes *pe/ppe* genes. Resulting backbone was recircularized. |
| 5. | pMV *esx-5<sub>Mxe</sub> ΔesxM/N* | pMV *esx-5<sub>Mxe</sub>* | XbaI, SnaBI | 7, 8<br>9, 10 | 1356 bp<br>1061 bp | Restricted backbone and the two PCR products were ligated in InFusion HD. |
| 6. | pMV *esx-5<sub>Mxe</sub> Δsubstr.* | pMV *esx-5<sub>Mxe</sub>* | AflIII, SnaBI | 11, 10 | 1082 bp | Restricted backbone and the PCR product were ligated with InFusion HD. |
| 7. | pMV *esx-5<sub>Mxe</sub> ΔespG<sub>5</sub>* | pMV *esx-5<sub>Mxe</sub>* | XbaI, SnaBI | 12, 13 | 2053 bp | Restricted backbone and the PCR product were ligated with InFusion HD. |
| 8. | pMV *ESX-5xen ΔeccD<sub>5</sub>* | pMV *esx-5<sub>Mxe</sub>* | SnaBI, NdeI |  |  | Restriction removes *eccD<sub>5</sub>*. Resulting backbone was recircularized. |
| 10. | pMV *ESX-5xen ΔeccE<sub>5</sub>* | pMV *esx-5<sub>Mxe</sub>* | NdeI, HindIII | 14, 15<br>16, 17 | 2068 bp<br>2607 bp | Restricted backbone and the two PCR products were ligated with InFusion HD. |
| 11. | pMV *ESX-5xen ΔEccA<sub>5</sub>* | pMV *esx-5<sub>Mxe</sub>* | MluI, HindIII | 18, 19 | 1129 bp | Restricted backbone and the PCR product were ligated with InFusion HD. |
| 12. | pMV *ESX-5xen MC* | pMV *esx-5<sub>Mxe</sub> ΔEccA<sub>5</sub>* | AflIII, SnaBI |  |  | Restriction removes all substrate genes and *espG<sub>5</sub>*. Resulting backbone was recircularized. |
| 13. | pSMT3 *pe/ppe* whole | pSMT3 empty |  | 20, 21 | 4583 bp | Backbone and PCR product were ligated with InfusionHD. |
| 14. | pSMT3 *pe/ppe Δpe/ppe<sub>1</sub>* | pSMT3 *pe/ppe* whole | NheI |  |  | Restriction removes 1st *pe/ppe* genes. Resulting backbone was recircularized. |
| 15. | pSMT3 *pe/ppe Δmethyltransf.* | pSMT3 *pe/ppe* whole | NheI, BamHI |  |  | Partial restriction to delete methyltransferase. Resulting backbone was recircularized. |
| 16. | pSMT3 *pe/ppe Δpe/ppe<sub>2</sub>* | pSMT3 *pe/ppe* whole | BamHI |  |  | Restriction remove 2nd *pe/ppe* genes. Resulting backbone was recircularized. |
| 17. | pMV *esx-5<sub>Mxe</sub> ppe<sub>1</sub> FLAG N-term* | pMV *esx-5<sub>Mxe</sub>* | AflIII, SnaBI | 22, 30<br>31, 25 | 688 bp<br>6035 bp | Restricted backbone and the two PCR products were ligated with InFusion HD. |
| 18. | pMV *esx-5<sub>Mxe</sub> ppe<sub>1</sub> FLAG C-term* | pMV *esx-5<sub>Mxe</sub>* | AflIII, SnaBI | 22, 23<br>24, 25 | 1934 bp<br>4769 bp | Restricted backbone and the two PCR products were ligated with InFusion HD. |
| 19. | pMV *esx-5<sub>Mxe</sub> ppe<sub>2</sub> Strep N-term* | pMV *esx-5<sub>Mxe</sub>* | AflIII, SnaBI | 22, 32<br>33, 25 | 3716 bp<br>3005 bp | Restricted backbone and the two PCR products were ligated with InFusion HD. |
| 20. | pMV *esx-5<sub>Mxe</sub> ppe<sub>2</sub> Strep C-term* | pMV *esx-5<sub>Mxe</sub>* | AflIII, SnaBI | 22, 26<br>27, 25 | 4910 bp<br>1802 bp | Restricted backbone and the two PCR products were ligated with InFusion HD. |
| 21. | pMV *esx-5<sub>Mxe</sub> esxN-HA* | pMV *esx-5<sub>Mxe</sub>* | AflIII, SnaBI | 22, 28<br>29, 25 | 5621 bp<br>1802 bp | Restricted backbone and the two PCR products were ligated with InFusion HD. |
| 22. | pMV *esx-5<sub>Mxe</sub> ppe<sub>1</sub> FLAG N-term, ppe<sub>2</sub> Strep C-term, esxN-HA, no Strep on eccC<sub>5</sub>.* | pMV *esx-5<sub>Mxe</sub> ppe<sub>1</sub> FLAG N-term* | AflIII, SnaBI | 22, 26 | 4910 bp | Restricted backbone and the two PCR products were ligated with InFusion HD into an intermediate plasmid. |
|  |  | pMV *esx-5<sub>Mxe</sub> esxN-HA* | AflIII, SnaBI | 27, 25 | 1802 bp |  |
|  |  | Intermediate plasmid | DraI, AflIII | 34, 35 |  | Restricted intermediate plasmid was religated with InFusion HD with the overlapping primers to swap the TwinStrep at the C-terminus of *eccC<sub>5</sub>* with a stop codon. |
| 23. | pMV *esx-5<sub>Mxe</sub> ppe<sub>1</sub> FLAG N-term, ppe<sub>2</sub> Strep C-term, esxN-HA, ΔeccC<sub>5</sub>* | pMV *esx-5<sub>Mxe</sub> ppe<sub>1</sub> FLAG N-term, ppe<sub>2</sub> Strep C-term, esxN-HA, No Strep on eccC<sub>5</sub>.* | PacI, AflIII | 5, 6 | 2174 bp | Restricted backbone and the PCR product were ligated with InFusion HD. |

